# Dorsal Periaqueductal gray ensembles represent approach and avoidance states

**DOI:** 10.1101/2020.11.19.389486

**Authors:** Fernando MCV Reis, Johannes Y Lee, Sandra Maesta-Pereira, Peter J Schuette, Meghmik Chakerian, Jinhan Liu, Mimi Q La-Vu, Brooke C Tobias, Newton S Canteras, Jonathan C Kao, Avishek Adhikari

## Abstract

Animals must balance needs to approach threats for risk-assessment and to avoid danger. The dorsal periaqueductal gray (dPAG) controls defensive behaviors, but it is unknown how it represents states associated with threat approach and avoidance. We identified a dPAG threat-avoidance ensemble in mice that showed higher activity far from threats such as the open arms of the elevated plus maze and a live predator. These cells were also more active during threat-avoidance behaviors such as escape and freezing, even though these behaviors have antagonistic motor output. Conversely, the threat-approach ensemble was more active during risk-assessment behaviors and near threats. Furthermore, unsupervised methods showed approach/avoidance states were encoded with shared activity patterns across threats. Lastly, the relative number of cells in each ensemble predicted threat-avoidance across mice. Thus, dPAG ensembles dynamically encode threat approach and avoidance states, providing a flexible mechanism to balance risk-assessment and danger avoidance.

## Introduction

Behavioral variables and emotional states are thought to be represented in neural activity (Anderson and Adolphs, 2014). Such representations must be specific enough to differentiate across behaviors, yet general enough to maintain functional cohesion across diverse threatening situations (Grundemann et al., 2019). A large body of evidence has shown that defensive behaviors related to threat exposure are represented in dorsal periaqueductal gray (dPAG) activity (Deng et al., 2016; Evans et al., 2018; Watson et al., 2016), as dPAG activity correlates with escape and freeze. Additionally, dPAG optogenetic and electrical stimulation induce these behaviors, as well as aversion (Brandao et al., 1982; Carvalho et al., 2015; Carvalho et al., 2018; Deng et al., 2016; Tovote et al., 2016). Furthermore, pharmacological manipulations of dPAG activity impact open arm exploration in the elevated plus maze (EPM), a traditional measure of rodent anxiety (Fogaca et al., 2012). Lastly, PAG activity in humans correlates positively with threat imminence (Mobbs et al., 2007; Mobbs et al., 2010). These reports show the dPAG is a central node orchestrating defensive behaviors.

However, it is unknown how the dPAG represents moment-to-moment changes in brain states during threat exposure. The two main behavioral states observed during exposure to threats are approach and avoidance (Stankowich, 2019). In the approach state, animals voluntarily go near the threat and perform risk-assessment behaviors. In this state, the exploratory risk-evaluation drive is stronger than the motivation to avoid danger. By contrast, in the avoidance state, animals perceive high risk, and thus attempt to minimize exposure to danger by escaping, freezing and maintaining distance to the threat. No reports to date have investigated whether the dPAG consistently encodes approach and avoidance states across distinct threats.

Key questions regarding the neural representation of approach and avoidance states remain unanswered. Do dPAG cells respond uniformly to transitions between higher and lower threat imminence? What is the overlap between the dPAG encoding of two completely distinct threats?

Does dPAG neuronal activity encode moment-to-moment changes regarding defensive approach and avoidance states? Addressing these questions would require population-level analysis of dPAG cells recorded longitudinally across threat modalities. Here, we report experimental data and analyses that directly address these questions.

## Results and Discussion

We performed microendoscopic calcium imaging of dPAG neurons expressing GCaMP6s (Figure 1A and Figure 1-figure supplement 1) (Cai et al., 2016) during EPM and rat exposure assays. During EPM test, we recorded 107 ± 19 cells per mouse (n = 8 mice; 857 cells were imaged; see Methods). As expected, mice spent more time in the closed arms of the EPM (Figure 1B). They also displayed exploratory risk-assessment head dips over the edges of the open arms (Figure 1B-C). During EPM exploration cells often showed preferential activity either in closed or open arms (Figure 1D-E). To identify EPM arm-type modulated neurons, we defined an “arm score metric” ranging from −1 to +1, in which the +1 indicates that cell activity in the open arm is greater than activity in the closed arm, and vice-versa. The arm score distribution in the observed data is wider than expected by chance, indicating that dPAG cells show robust preference for EPM arm-types (Figure 1F left panel). We defined neurons as belonging to one of the ensembles if activity in each arm-type was significantly greater than the pooled activity in the opposite arm type (Figure 1F right panel, see Methods). The results showed that cells fired similarly in arms of the same type, as firing in one open arm was highly correlated to activity in the other open arm (Figure 1G, top panel). Conversely, firing rates in closed and open arms were negatively correlated (Figure 1G, bottom panel).

**Figure 1.**
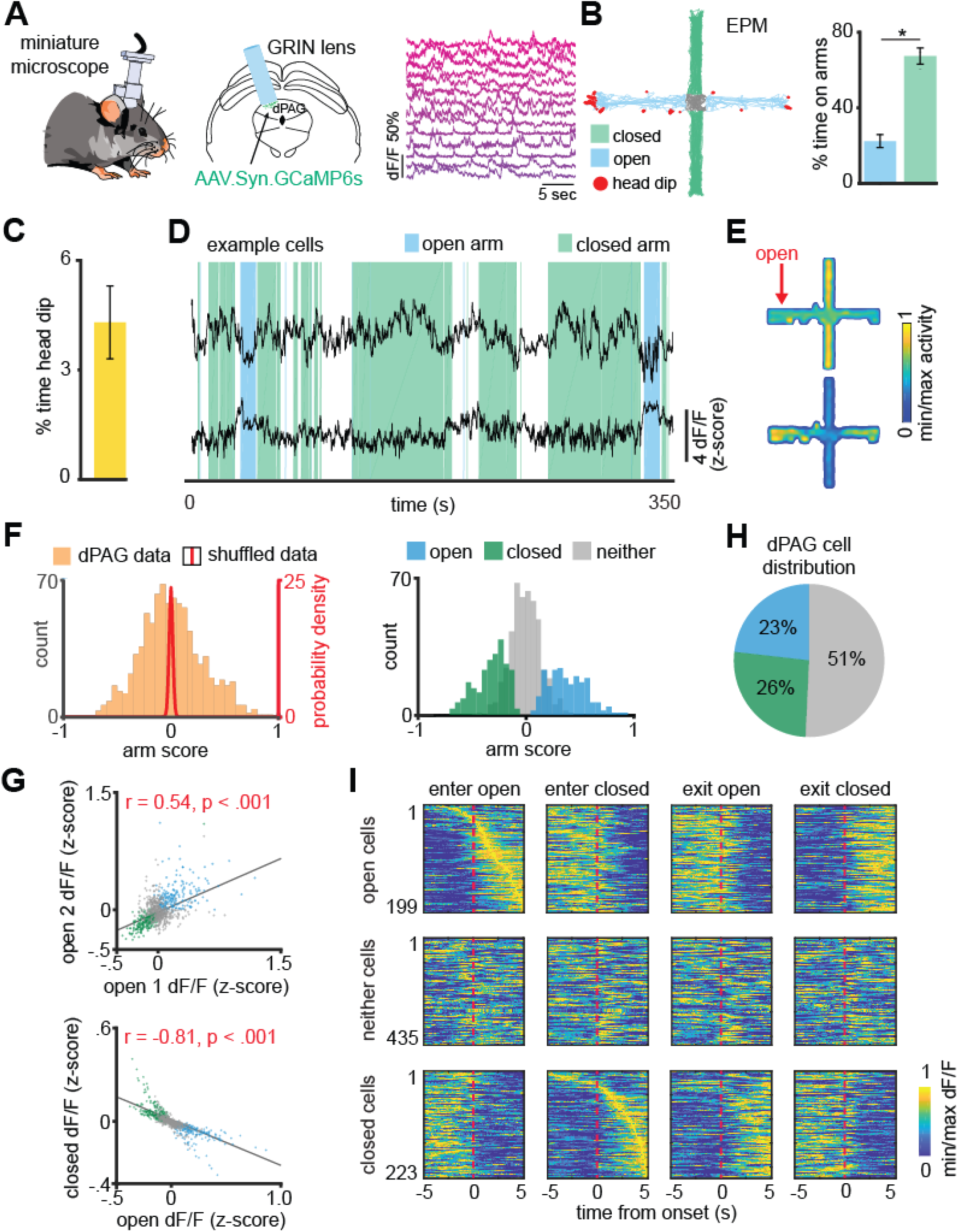
DPAG neuronal ensembles encode arm-type in the elevated plus maze. (**A**) GRIN lens implantation, virus expression strategy and example Ca2+ signals of neurons the dorsal periaqueductal gray (dPAG). (**B**) Example mouse exploration path recorded in the EPM. Mice spent significantly more time in the closed arms compared to the open arms (Data are represented as mean ± SEM; *W* = 0, *p* = 0.012, Wilcoxon Signed Rank Test, *n* = 8 mice;). (**C**) Mean percentage of total time in which mice engaged in head dips. (*n* = 8 mice). (**D**) dPAG dF/F traces from the same mouse that display preferential firing in the closed (upper trace) and open (lower trace) arms of the EPM (open and closed arm-preferring cells). Epochs corresponding to exploration of the closed and open arms are shown respectively as green and blue shaded areas. (**E**) Activity heat maps for corresponding example neurons shown in (**D**). (**F**) The arm preference score was calculated for each neuron (orange bars; see Methods), as was the distribution of arm preference scores for shuffled data (red line). Bars show the distribution of arm preference scores for open, closed, and neither cells. (*n* = 857 cells). (**G**) Scatterplots showing correlations between neural activity across the two open arms (top) and between open and closed arms of the EPM (bottom). Each point represents one cell (*n* = 857 cells*, r* = Pearson’s correlation coefficient). (**H**) Pie chart shows the percent of all recorded neurons that were classified as open, closed or neither cells. (*n* = 857 cells). (**I**) For each subplot, each row depicts the mean normalized activity of an open, closed, or neither arm-preferring cell during behavior-aligned arm transitions. (*n* = 857 cells).

Based on the distribution of cells per arm score, roughly half of the dPAG neurons were classified as arm-modulated cells (49%, with 26% closed- and 23% open-modulated cells) (Figure 1H) which suggests these ensembles are functionally relevant dPAG populations. During transitions between arms, we identified opposite changes in activity levels of these two major, non-overlapping populations of dPAG neurons. (Figure 1I and Figure 2A-B). For example, the closed arm-activated ensemble showed a decrease in activity when mice traversed from a closed arm to an open arm. Moreover, open and closed cells showed increased and decreased activity, respectively, during exploratory head dip behavior (Figure 2C). Importantly, dPAG ensembles did not display strong correlations with speed, suggesting these findings are not driven by variations in velocity (Figure 2D). If EPM arm-type is prominently represented in dPAG activity, then it may be possible to use dPAG activation patterns to differentiate mouse location in the EPM. Indeed, upon training support vector machine (SVM) decoders on dPAG activity, we obtained significantly higher than chance performance in identifying if the mouse was in an open or closed arm (Figure 2E, see Methods).

**Figure 2.**
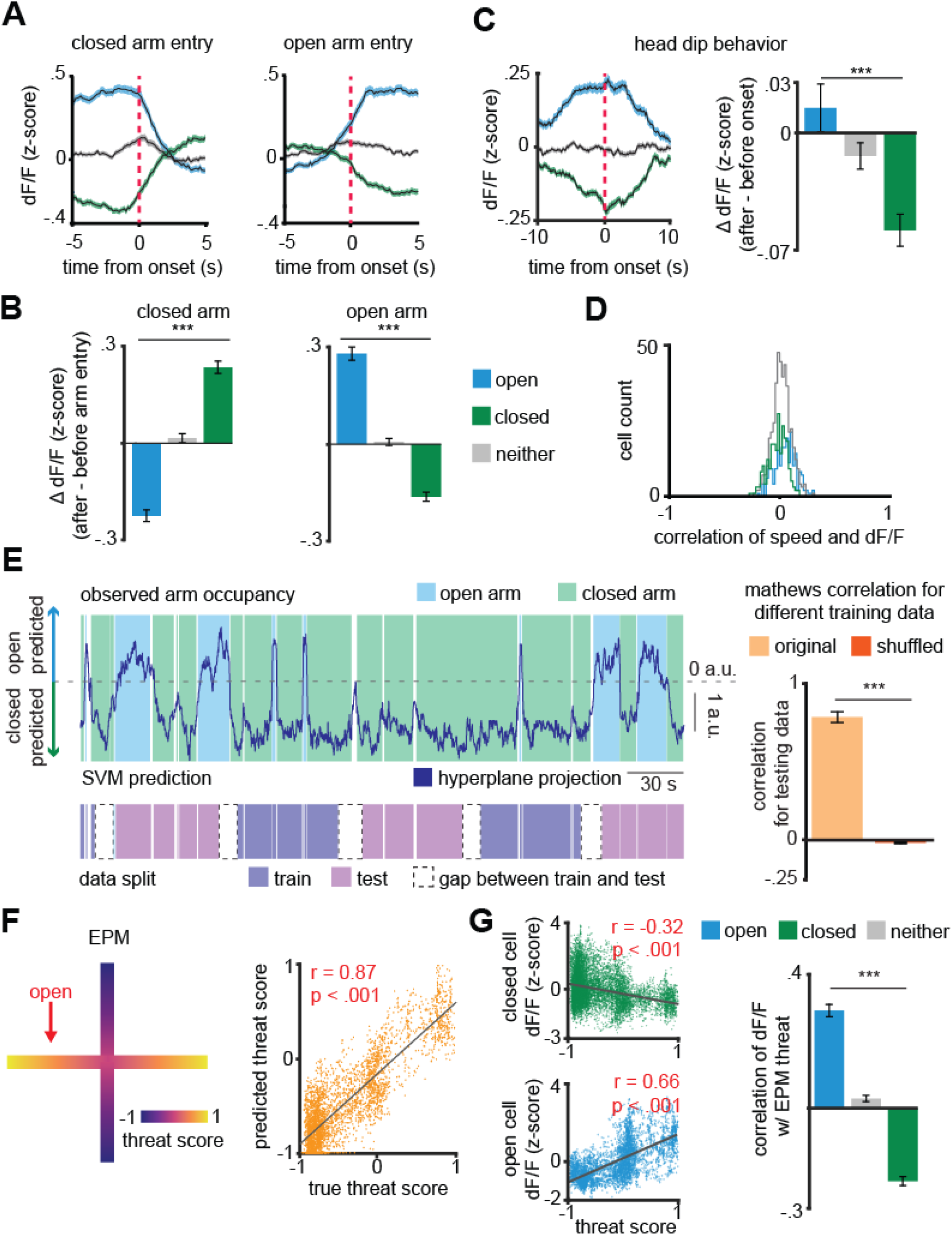
DPAG population activity predicts EPM exploration. (**A**) Traces show the mean z-scored activity (+/− 1 SEM) of all open, closed, and neither cells, behavior-aligned to arm transitions (respectively the blue, green and gray traces). (**B**) Bars depict the change in z-scored dF/F for entries to closed left) and open (right) arms, separately for open, closed and neither cells. (Data are represented as mean ± SEM; both closed and open arm; *n* = 199 open cells, *n* = 435 neither cells, *n* = 223 closed cells; closed arm *U* = 14.49, \*\*\**p* < 0.001, open arm *U* = 14.05, *p* < 0.001, Wilcoxon Rank Sum Test, *n* = 8 mice). (**C**) Average activity traces for open, closed, and neither cells relative to onset of head dips in the EPM and quantification of changes in activity for all cell types (0-2.5 seconds after minus 2.5-5.0 seconds before head dip onset) (Data are represented as mean ± SEM; *n* of cells same as (G); *U* = 4.53, \*\*\**p* < 0.001, Wilcoxon Rank Sum test). (**D**) Histograms depict the distribution of the Pearson correlation of dF/F with speed for each cell type in the EPM. (**E**) Prediction of arm-type mouse position in the EPM from neural data using a linear support vector machine (SVM). The blue and green areas represent the actual arm-type occupancy label (open and closed arm, respectively), and the black trace represents the prediction of arm location by the SVM hyperplane projection. If the trace was above 0 a.u., then that period was classified as open arm exploration, otherwise, it was classified as closed arm occupancy. The pink and purple represent the data split (training and testing data, respectively). The matthews correlation coefficient for real and permuted shuffled training data are shown to the right (mean +/− 1 s.e.m.; n = (8, 800), *U* = 4.87, p < .001, Wilcoxon Rank Sum Test). (**F**) (left) Example of EPM threat score where 1 and −1 correspond, respectively, to the extreme end of the open and closed arms and (right) prediction of labeled EPM threat from dPAG neural data for an example mouse (scatterplot displays testing data that was not used for training, *r* = Pearson’s correlation coefficient). (**G**) (left) Correlation of example closed and open cell activity with threat score and (right) mean correlation of dF/F with EPM threat (Data are represented as mean ± SEM; *n* = 64 open cells, *n* = 166 neither cells, *n* = 87 closed cells; EPM *U* = 10.48 Wilcoxon Rank Sum Test, \*\*\**p* < 0.001, *r* = Pearson’s correlation coefficient).

Next, we investigated whether dPAG activity could also predict specific mouse positions within the arms. We defined a “threat score” as linearly varying between −1 and +1 (at closed and open arms extremities, respectively), assigning zero to the center of the maze (Figure 2F, see Methods). We then fitted a linear regression on the threat score using dPAG cell activity. Interestingly, the model showed significantly higher than chance performance in predicting threat score, suggesting that the identified dPAG ensembles may not only encode arm-type, but rather a risk perception and threat-exposure gradient (Figure 2F-G and Figure 2-figure supplement 1).

To investigate whether dPAG population coding of risk perception generalizes to exploratory behavior across different threatening contexts, we recorded the same dPAG neurons during exposure to a live predator (Figure 3A-B and Figure 3–figure supplement 1). Mice were allowed to freely explore a context, a long chamber, in the presence of a rat, which was tethered with a harness to one end of the chamber (see Methods). All behavioral data from synchronized videos underwent automated behavior scoring (Mathis et al., 2018). Mice spent most of the trial away from the rat (Figure 3A), indicating aversion from perceived threat. Consistent with previous threat imminence theories and the array of defensive behaviors evoked in the presence of a predator (Blanchard et al., 2011; McNaughton and Corr, 2004; Perusini and Fanselow, 2015; Stankowich, 2019), mice presented defensive strategy repertoires composed of approach and avoidance-related behaviors (i.e. escape and freeze) (Figure 3A). Notably, average dPAG activity increased with rat proximity, rat movement onset, escape, and decreased during freezing and approach (Figure 3C-E). Importantly, mice displayed no signs of aversion nor differences in dPAG activity with proximity to a toy rat (Figure 3–figure supplement 2).

**Figure 3.**
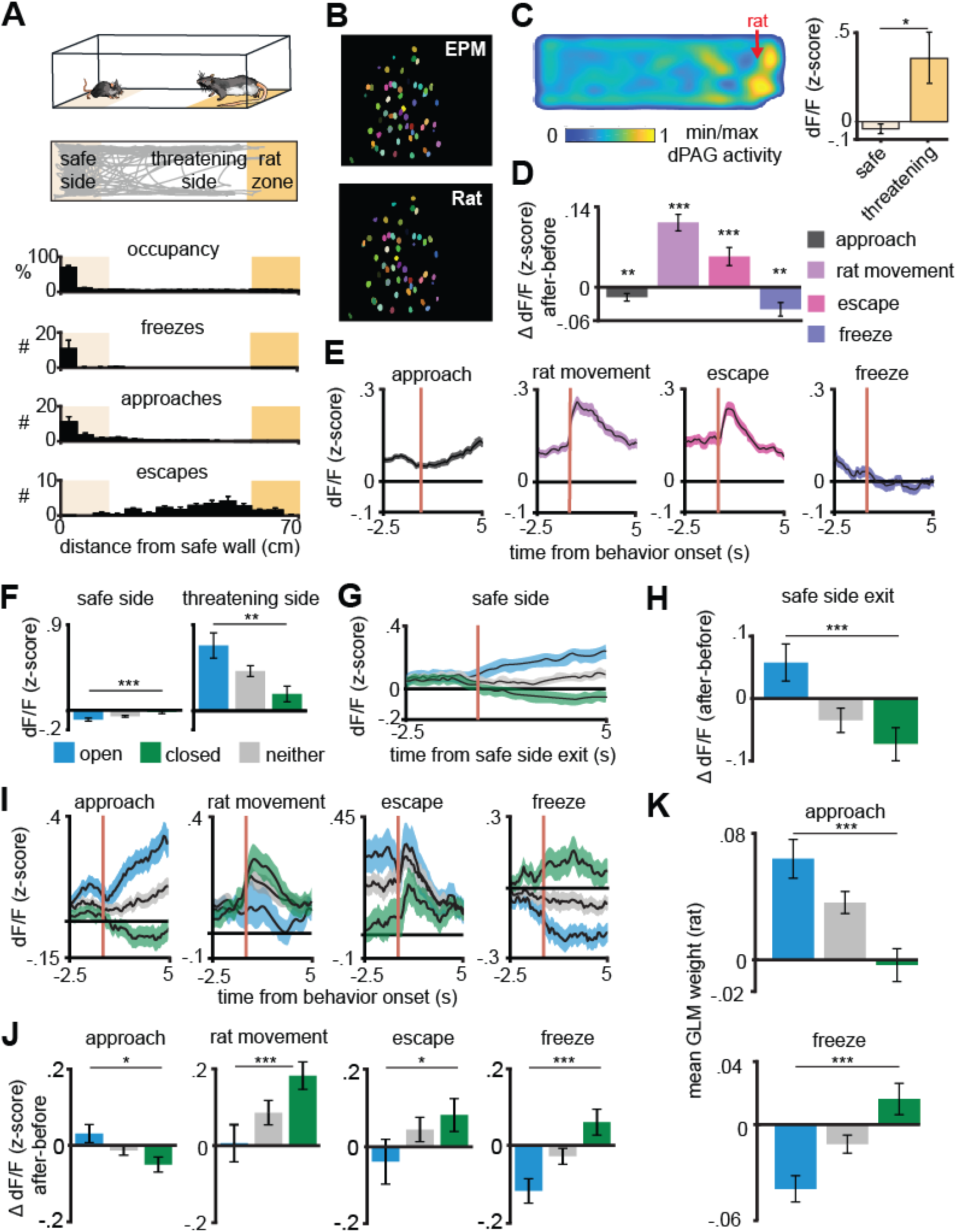
Arm-specific ensembles maintain functions across threatening situations. (**A**) Illustration of the rat exposure assay (top) and example track (bottom), with labels depicting the area to which the rat is confined (rat zone) as well as areas near to and far from the rat (safe side). In all figures depicting this assay the rat area will be shown to the right. (**B**) Example imaging field of view with dPAG cells co-registered between EPM and rat exposure sessions. (**C**) Heatmap depicts the mean z-scored dF/F at each position of the rat exposure assay (*n* = 713 cells, *n* = 7 mice). (**D**) Change in dF/F (0-2.5 seconds after minus 0-2.5 seconds before) activity for all dPAG cells for behaviors in the rat exposure assay (Data are represented as mean ± SEM; *n* of cells for approach, stretch, escape = 714, *n* of cells for freeze = 640; approach *t* = −2.65, \*\**p* = 0.008, stretch *t* = 4.92, \*\*\**p* < 0.001, escape *t* = 3.39, ***p < 0.001, freeze *t* = −3.23, \*\**p* = 0.0012, one-sample *t*-test). (**E**) Traces show the mean z-scored activity of all cells (+/− 1 s.e.m.), aligned to onset of various behaviors (onset is indicated by the red vertical line) in the rat exposure assay (*n* of cells same as D). (**F**) Bars depict the mean z-scored dF/F of cells on the safe side and threatening side of the enclosure (Data are represented as mean ± SEM; *n* = 64 open cells, *n* = 166 neither cells, *n* = 87 closed cells; *n* = 7 mice, safe *U* = −3.82, \*\*\**p* = 0.0001, threatening *U* = 3.05, \*\**p* = 0.002, Wilcoxon Rank Sum Test). (**G**) Traces show the mean z-scored activity of open, closed, and neither cells (+/− 1 s.e.m.), aligned to exit of the safe side of the enclosure (far from the rat). (**H**) Bars show the mean change in z-scored dF/F (0-2.5 seconds after minus 0-2.5 seconds before) aligned to safe side exit for open, closed, and neither cells. (Data are represented as mean ± SEM; *n* = 64 open cells, *n* = 166 neither cells, *n* = 87 closed cells; *U* = 3.36, \*\*\**p* = 0.0008, Wilcoxon Rank Sum Test). (**I**) Traces show the mean z-scored activity of open, closed and neither cells (+/− 1 SEM), aligned to behaviors in the rat exposure assay (approach, escape: *n* = 64 open cells, *n* = 87 closed cells, *n* = 166 neither cells; stretch: *n* = 50 open cells, *n* = 64 closed cells, *n* = 127 neither cells; freeze: *n* = 39 open cells, *n* = 56 closed cells, *n* = 117 neither cells). Onset of behaviors is indicated by a red vertical line. (**J**) Bars depict the change in z-scored dF/F (0-2.5 seconds after minus 0-2.5 seconds before) for behaviors in the rat exposure assay, separately for open, closed, and neither cells (Data are represented as mean ± SEM; *n* of cells same as (**I**); approach *U* = 2.45, \**p* = 0.014, stretch *U* = 0.25, *p* = .81, escape *U* = −2.16, \**p* = .03, *U* = −3.80, \*\*\**p* = 0.0001, Wilcoxon Rank Sum Test). (**K**) A generalized linear model (GLM) to predict single cell activity was constructed using approach, stretch, escape and freeze behaviors as variables. Bar plots show average GLM weights for approach and freeze for open, closed and neither cells. (Data are represented as mean ± SEM; *n* = 62 open cells, *n* = 155 neither cells, *n* = 83 closed cells; approach *U* = 4.17, \*\*\**p* < 0.001, *p* = .11, freeze *U* = 3.02, \*\*\**p* < 0.001, Wilcoxon Rank Sum Test).

We then explored whether the activity of dPAG closed and open cell ensembles identified in the EPM also represent risk-evaluation in the rat assay. A positive result would show that the ensembles are likely responding not only to the original sensory biases (i.e. closed and open arms features), but potentially representing behavioral states that generalize across threats. Intriguingly, open arm cells were more active near the rat, while closed arm cells displayed higher activity far from the rat (Figure 3F-H and Figure 3-figure supplement 3). Closed cells also showed increased activity following onset of both escape and freeze, despite these behaviors having opposite motor outputs (Figure 3I-K). Additionally, even though freezing and approach onset occur similarly far from the rat (Figure 3A), open and closed ensembles showed opposite activity patterns and different generalized linear model weights (Figure 3I-K). Rat movement onset likely constitutes a threat signal and switches states from approach to avoidance, as predator movement is indicative of increased threat imminence. Indeed, rat movement is a significant predictor variable for less frequent approach and more occurrences of threat avoidance-related behaviors, such as escape and freezing (Figure 3-figure supplement 4). Accordingly, during rat movement onset, the threat-avoidance related closed-arm ensembles displayed higher activity (Figure 3I-J). Notably, neither of the ensembles consistently resembled the overall average dPAG activity during rat exposure (Figure 3E), as each ensemble had its own functional profile (Figure 3I). Furthermore, dPAG cells also used shared patterns of neural activity across rat and EPM assays to represent threat imminence (measured as distance to threat) (Figure 3-figure supplement 5, see methods). These results suggest that dPAG neuronal activity can represent internal brain states using shared patterns of activity across different threats.

An approach-state is associated with open arm entries, head dips in the EPM and proximity to threat. Conversely, an avoidance-state would be expected far away from threats and during actions that decrease threat exposure, such as closed arm exploration, escape and freezing. Our results showed that closed cells were more active during higher distance from threat and threat avoidance-related behaviors, such as freezing and escaping, while open cells were more active during proximity to threats and exploratory head dips in the EPM (Figure 2 and Figure 3). The consistency of these results across behaviors and two different threat modalities indicate that dPAG closed and open cells were encoding threat avoidance and threat approach states, respectively.

To further investigate how dPAG cells use a shared representation to encode approach and avoidance states we developed an approach/avoidance score ranging between −1 and 1 (see Methods). The score gradually increases during approach to threat and during EPM head dips, reaching +1 when the mouse is adjacent to the rat or in the extreme end of the open arms. The score decreases when the mouse is retreating from threat and is assigned a value of −1 when the mouse is furthest from the rat, freezing, or in the extreme end of the closed arms (Figure 4A). To investigate how the approach/avoidance score is encoded in dPAG activity we used k-means clustering, an unsupervised approach, to group the data points into 10 clusters (Figure 4B, panel 1). We chose 10 clusters, as opposed to 2, since the neural data does not exclusively encode approach/avoidance states. We then calculated the approach/avoidance score for each of the 10 clusters (Figure 4B, panel 2). The clusters with the lowest and highest scores were classified as the ‘avoidance’ and the ‘approach’ cluster, respectively. Experimentally observed approach and avoidance clusters respectively had higher and lower scores than bootstrap distributions, showing that our k-means approach identified activity patterns that strongly encode the approach-avoidance score (Figure 4C). The approach and avoidance cluster centroids from one assay were then applied to the other assay (i.e., centroids were defined by training on EPM and applied on previously unseen data from the rat assay, or vice-versa). Cluster centroids defined from the training data in one assay were applied to the data from the other assay to assign cluster identity based on shortest euclidean distance to the centroid. For example, the points in the testing dataset that were closest to the avoidance centroid, previously defined by the training dataset, were assigned to the avoidance cluster (Figure 4B, panels 3-4). Scores for approach and avoidance clusters for shuffled data were used to create a bootstrap distribution. Lastly, we show that approach and avoidance clusters trained on one assay and applied on the other assay result in significantly different scores, despite the two assays having different geometries and distinct threat modalities (Figure 4D). Similar results were also obtained for k-means using a smaller number of clusters or employing a hidden Markov model, showing that the results in Figure 4 can be found using a range of different computational approaches (Figure 4-figure supplement 1). Importantly, these results were not found when computing approach and avoid clusters centroids defined on the EPM but applied to a control toy rat (Figure 4-figure supplement 2). These results indicate, using an unsupervised method, that approach and avoidance states are encoded using shared patterns of neural activity across assays. Importantly, dPAG activity reflects moment-to-moment changes in the behavioral states of the animal.

**Figure 4.**
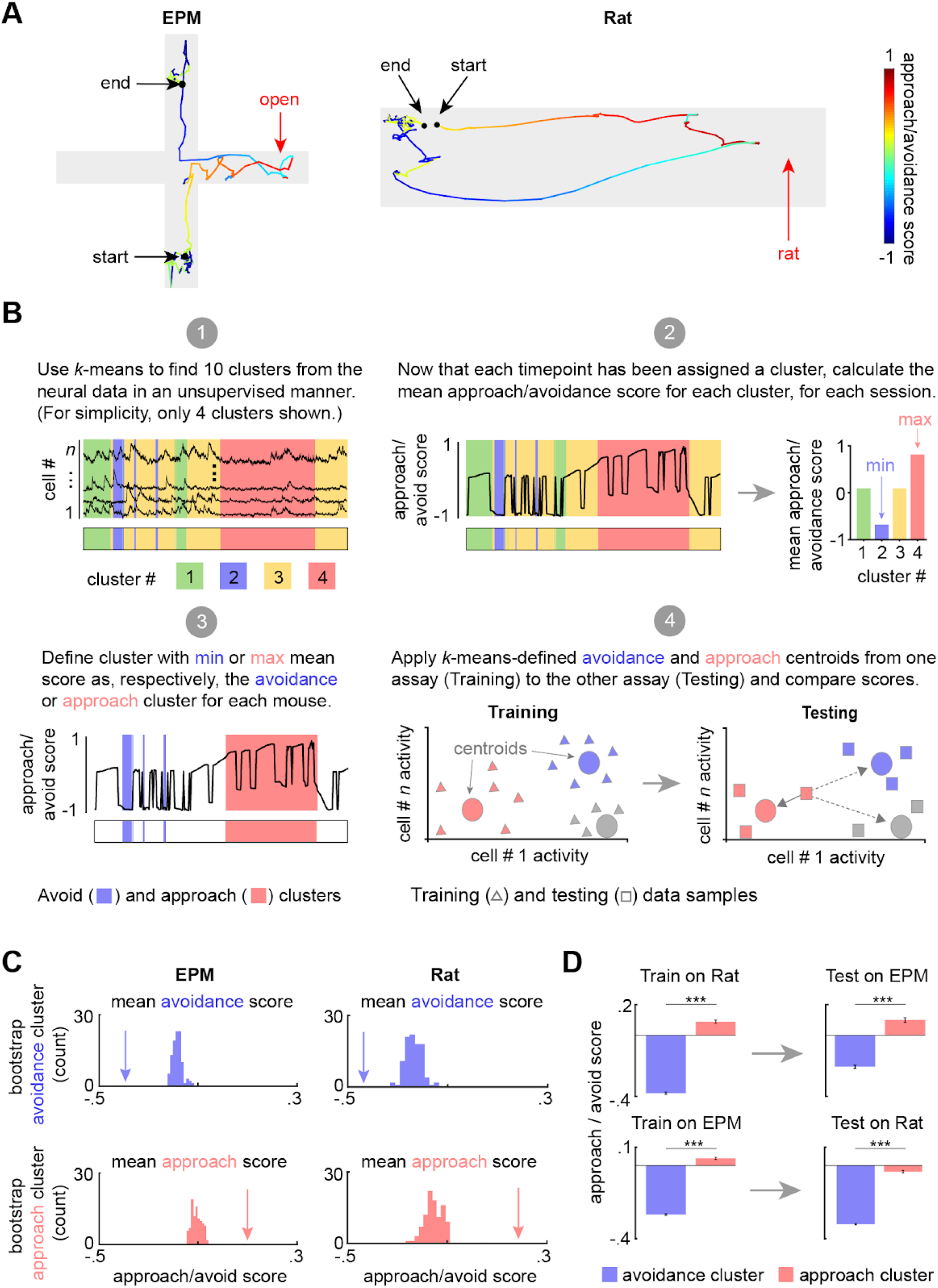
DPAG displays a shared neural representation of approach and avoidance states across the EPM and rat exposure assays. Example tracks in the EPM (left) and Rat exposure assay (right), color-coded by approach/avoidance score (see Methods). The approach score increased during movements towards the threat, reaching +1 when the animal reaches the end of the open arms or the rat. The score decreases during movement away from threat and reaches its minimum value of −1 when the mouse reaches the end of closed arms or the furthest point from the rat. This score was developed as a measure of approach/avoidance states. (**B**) Explanatory diagram depicting steps of the clustering analysis (see Methods). *K*-means (k = 10) was used to find clusters in the neural data in an unsupervised manner. (2) The mean approach/avoidance score was calculated for each cluster defined in step 1. (3) The ‘avoidance’ and ‘approach’ clusters were identified as those with, respectively, the minimum or maximum mean approach/avoid score calculated in step 2. (4) The approach and avoidance centroids defined in one assay were used to classify neural data from the other assay, based on the minimum euclidean distance for each sample (as depicted by solid arrow). (**C**) Arrow depicts the experimentally observed mean approach/avoidance score for avoidance and approach clusters across concatenated sessions (*n* = 7 mice). This mean was compared to a bootstrapped distribution of avoidance (top) or approach (bottom) cluster means, calculated by shuffling the neural data 100 times (EPM cells n=801, rat assay cells n=878; for all, *p* < 0.01). (**D**) (left) Bars depict the mean Rat and EPM approach/avoidance scores (+/− 1 s.e.m.) for approach and avoidance clusters across mice. (right) As described in Methods and 4B, these cluster centroid locations, trained on one assay, were then used to define approach and avoidance timepoints in the other assay. Bars depict the corresponding mean approach/avoidance score (+/− 1 s.e.m.) for this testing data (Train on EPM: avoidance cluster *n* = 3867, approach cluster *n* = 5027; Test on Rat: avoidance cluster *n* = 2445, approach cluster *n* = 2622; Train on Rat: avoidance cluster *n* = 4894, approach cluster *n* = 3222; Test on EPM: avoidance cluster *n* = 2624, approach cluster *n* = 1446 (n represents the number of time points, not cells); coregistered cells n=399; Wilcoxon ranked-sum test, *** p < 0.001).

Finally, we investigated if ensemble composition was related to threat avoidance traits across assays. Rat approach and open arm exploration was correlated across mice, indicating that these measures also reflected trait avoidance levels (Figure 4-figure supplement 3A). We then found that mice with a higher proportion of open cells in relation to closed cells displayed increased avoidance of open arms and rat (Figure 4-figure supplement 3B-C). These data indicate that in addition to encoding moment-to-moment changes on behavioral states, dPAG ensembles composition may integrate risk evaluation processes and influence individual mouse differences in threat avoidance traits.

Together, these results suggest that the dPAG neural population has a shared representation of risk perception across threatening circumstances. Individual neurons change their activity similarly to represent threat approach and avoidance states across assays. These findings expand on the oversimplified view of dPAG as a pre-motor output region and highlights it as a key node reflecting the internal brain states that prepare the organism to engage in approach or avoidance of threat.

## Acknowledgements

This work was supported by the National Institute for Mental Health (R00 MH106649 and R01 MH119089) (A.A.), the Brain and Behavior Research Foundation (Grants # 22663 and 27654 respectively to AA and FMCVR), the National Science Foundation (NSF-GRFP DGE-1650604, P.J.S.), the UCLA Affiliates fellowship (P.J.S.) and the Hellman Foundation (A.A.). Fundação de Amparo à Pesquisa do Estado de São Paulo (FAPESP), Research Grant #2014/05432-9, (NSC). FMCVR was supported with FAPESP grants #2015/23092-3 and #2017/08668-1. MQLV was supported by the Achievement Rewards for College Scientists Foundation and by the National Institute for Mental Health award F31 MH121050-01A1.

## Author contributions

FMCVR, NSC and AA conceived the project and designed the experiments. FMCVR, AA, JYL, SM and JCK wrote the paper. FMCVR, MC and SM performed the behavioral assays. JYL, PJS, SM and JL analyzed the data. MQL and BCT assisted in the rat assay. JCK and AA supervised the data analysis.

### Supplementary Figures

**Figure 1–figure supplement 1.**
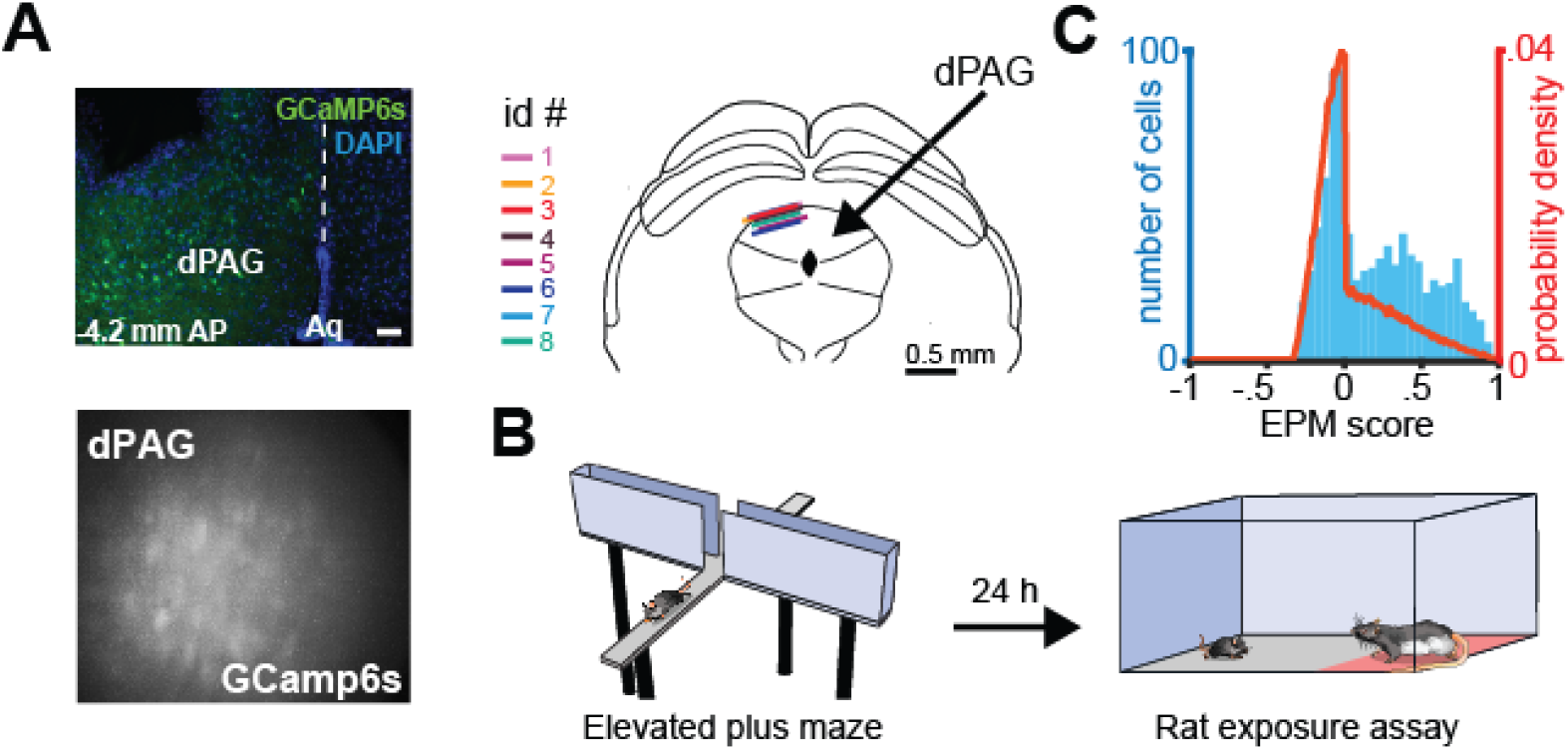
Deep brain imaging of dorsal periaqueductal gray neurons and distribution of EPM scores. (**A**) (top) Coronal section of the periaqueductal gray showing GCaMP6s expression and representative GRIN lens placement in the dorsal part of the periaqueductal gray (dPAG) (DAPI, blue; GCaMP6s, green). Position of the section relative to bregma is indicated in the lower left corner. Aq: aqueduct (Sylvius). Scale bar: 50um. (bottom) Maximum projection of the dPAG field of view in an example mouse. (top right) Anatomical scheme of GRIN lens front location of animals expressing GCaMP6s under the control of the Syn promoter in large populations of dPAG neurons (*n* = 8 mice). (**B**) Illustration and sequence of elevated plus maze (EPM) and rat exposure assays. (**C**) The EPM score (blue) differs significantly from distribution expected by chance (red line). (*n* = 857 cells; *U* = 15.62, *p* <0 .001, Wilcoxon Rank Sum Test). The EPM score is near +1 for cells that fire similarly in arms of the same type. For example, a cell that had very high firing rate in both open arms and very low rate on both closed arms would have a score near 1. A cell with high firing in both closed arms and low firing in both open arms also would have a score near 1. Thus, both open and closed-arm preferring cells would have positive scores. A cell firing uniformly in the environment would have a score near 0. A cell that fired differently in arms of the same type would have negative score, such as a cell that has high firing rate in one open arm and lower than mean firing rate in the other open arm. Note that this is a different metric from the arm score metric shown in Figure 1F.

**Figure 2–figure supplement 1.**
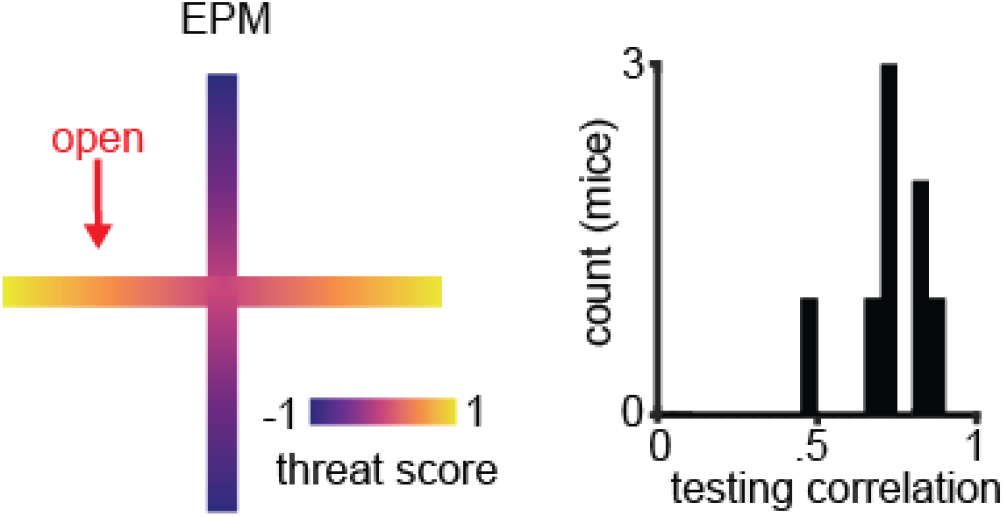
Validation of linear regression to predict threat score using dorsal periaqueductal gray activity. Example of EPM threat score where 1 and −1 correspond, respectively, to the extreme end of the open and closed arms. Histogram of testing correlations between labeled and predicted threat for individual mice is significantly greater than zero (*t* = 17.7, *p* < 0.001, 7 d.o.f., one-sample *t*-test, *n* = 8 mice).

**Figure 3–figure supplement 1.**
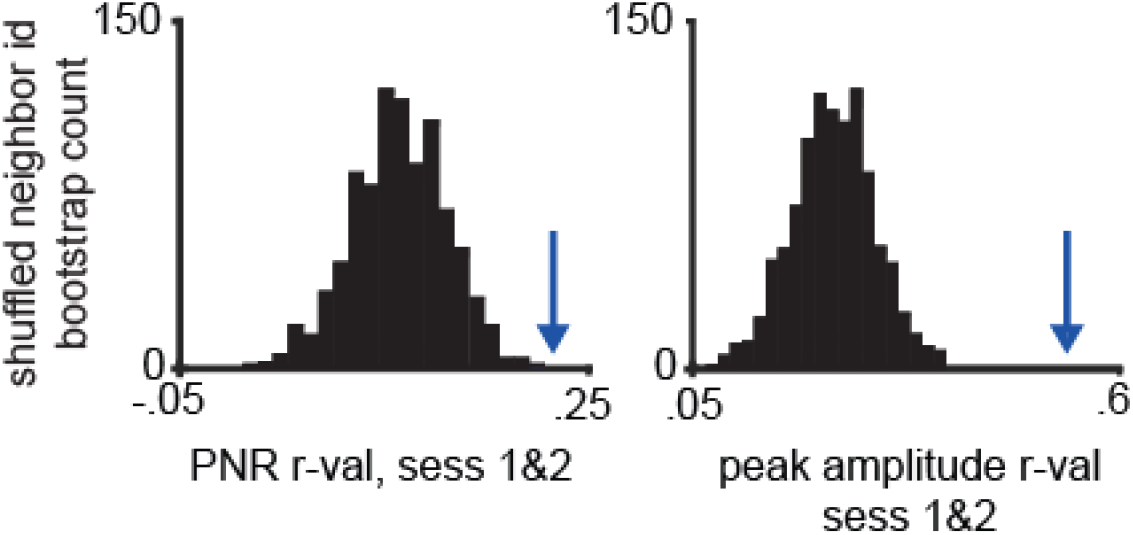
Validation of co-registration procedure. The peak-to-noise ratio (PNR) and mean peak amplitude correlation values were calculated for co-registered cells between rat and EPM assays. Cell identities were then shuffled within the ten nearest neighbors 1000 times, and the same correlation measures were calculated for each iteration. The resulting bootstrap distribution was compared to the actual peak-to-noise and mean peak amplitude values, indicated with arrow (*n*=462; *p*<.001, *n*=7 mice).

**Figure 3–figure supplement 2.**
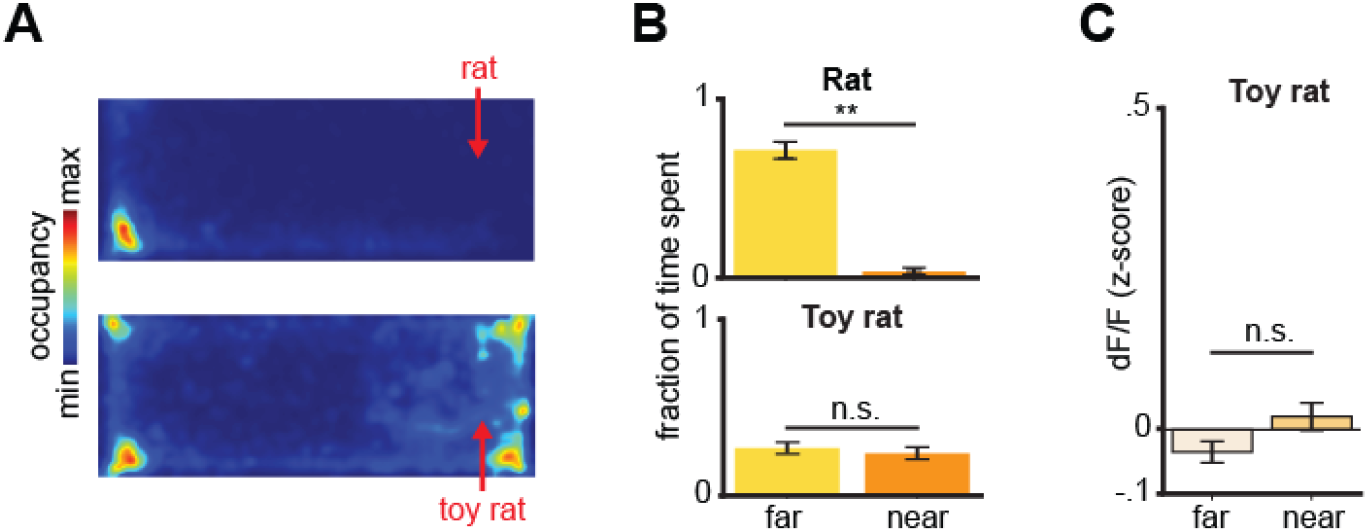
Behavioral and dPAG activity differences between rat and toy rat exposure. (**A**) Heatmaps depict spatial occupancy during rat (top) and toy rat exposure (bottom). (**B**) Time spent near and far from rat (top, *U* = 3.13, *p* = 0.0017, Wilcoxon Rank Sum Test) and toy rat (bottom, *U* = 0.70, *p* > 0.05, Wilcoxon Rank Sum Test). (**C**) Average z-scored dPAG activity far and near to a toy rat (*U*=1.85, *p* > 0.05, Wilcoxon Rank Sum Test). (A-C, *n*=7 mice).

**Figure 3–figure supplement 3.**
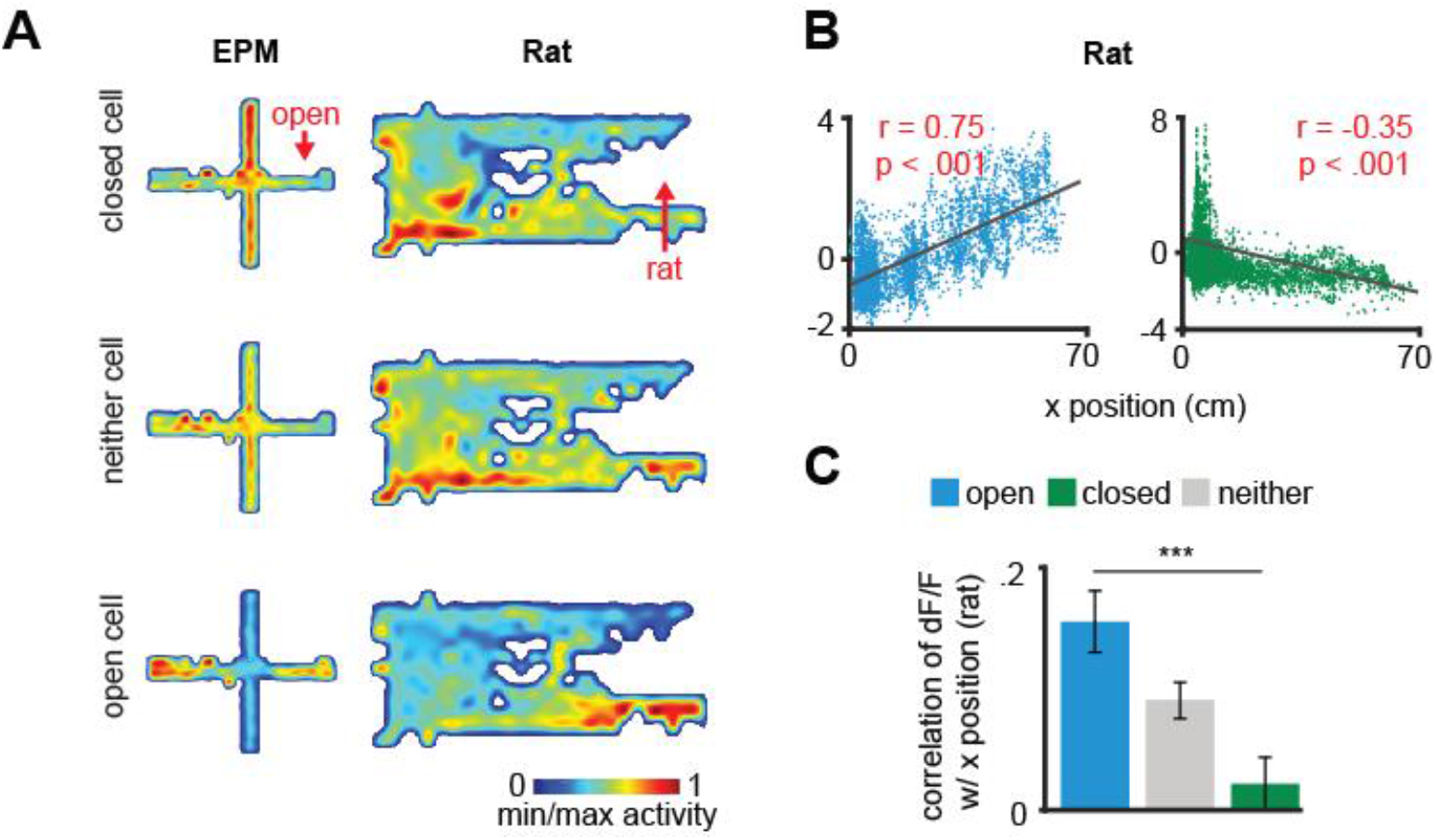
Correlation of dorsal periaqueductal gray cell ensembles with distance to a rat. (**A**) Heatmaps show the normalized activity of example open, closed, and neither cells across both the EPM and rat exposure assays. Note that the open arm cell was more active near the rat while the closed cell was more active far from the rat. Correlation of example open and closed cell activity with distance from the safe wall (rat exposure). Higher x position corresponds to locations more near the rat. (**C**) Mean correlation of dF/F with x position in the rat assay (Data are represented as mean ± SEM; *n* = 64 open cells, *n* = 166 neither cells, *n* = 87 closed cells; *U* = 3.89, \*\*\**p* < 0.001, Wilcoxon Rank Sum Test, *n* = 7 mice, *r* = Pearson’s correlation coefficient).

**Figure 3–figure supplement 4.**
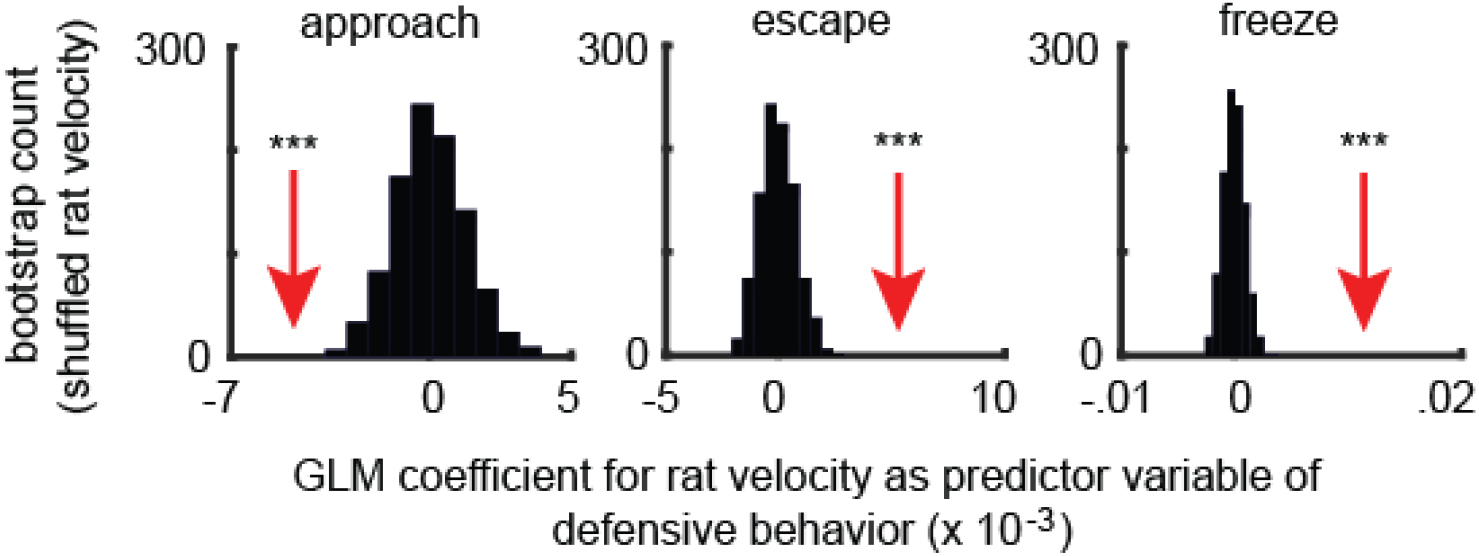
Increased rat velocity predicts lower approach to rat and higher threat avoidance related behaviors such as escape and threat. Separate generalized linear models (GLMs) were fit with rat velocity as the predictor variable and one of the binarized mouse behaviors: either approach (left), escape (middle), or freeze (right), as the response variable. The red arrow depicts the actual GLM coefficient for rat velocity, given each mouse behavior, while the histogram depicts the bootstrapped distribution of rat velocity coefficient values for shuffled timepoints. Compared to this distribution, rat velocity shows a significantly negative coefficient for approach and significantly positive coefficients for escape and freeze. These data show that higher rat velocity predicts decreased occurrence of approach and increased frequency of escape and freezing. (*** p < 0.001, *n* = 7 mice).

**Figure 3–figure supplement 5.**
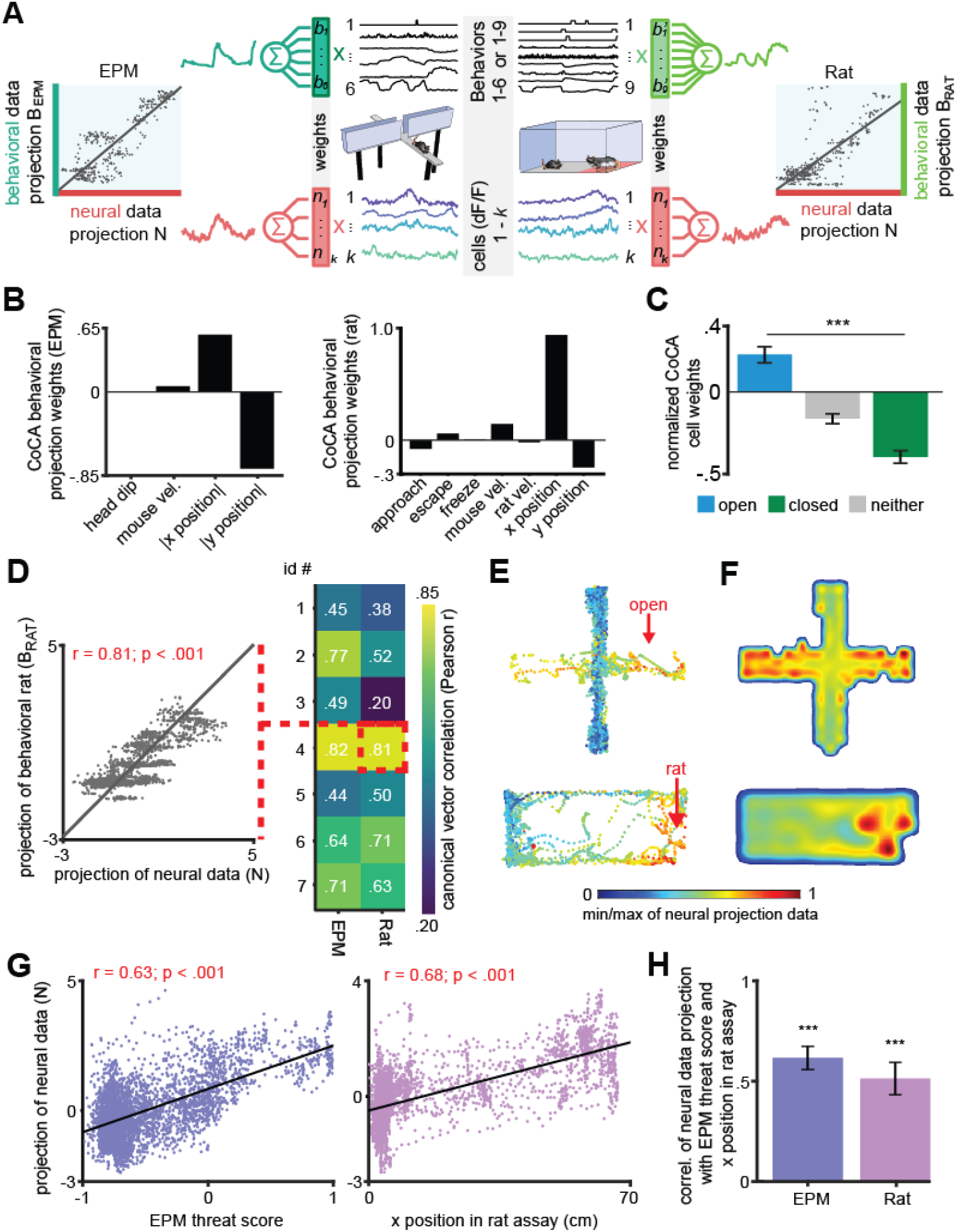
DPAG displays a shared neural representation of risk imminence across the EPM and rat exposure assays. (**A**) Constrained Correlation Analysis (CoCA) reveals correlated encoding of behaviors and neural activity consistent across EPM and rat exposure assays. Linear projections of behaviors (top) correlate with projections of neural activity (bottom). Weights were optimized to maximize the correlation between neural and behavioral projections. Weight selection was constrained in the following way: weights for behavioral variable weights for each assay had to be conserved across mice, whereas neural projector weights had to be fixed across the EPM and rat assays for each cell. The behavioral variables used are listed in the x-axis of (B). Colors indicate consistent cells and projector weights. (**B**) Weights of CoCA behavioral projector variables for the EPM (top) and rat exposure (bottom) assays showing the relative importance of each variable in each assay. All variables were normalized to unit variance before training and testing, with the exception of |x| and |y|, which were scaled to the range [0, 1]. (**C**) CoCA neural projection weights normalized to the range [−1, 1], mean +/− 1 SEM. (Data are represented as mean ± SEM; *n* = 64 open cells, *n* = 166 neither cells, *n* = 87 closed cells; *U* = 8.02, \*\*\**p* < 0.001, Wilcoxon Rank Sum Test, *n* = 7 mice). (**D**) (left) Example correlation of CoCA projection of behavioral data with projection of neural data for testing data (mouse 4, rat exposure). Each point is one time point of data. (right) Correlation values of CoCA projection of behavioral data with projection of neural data for testing data for each mouse in each assay (*p* < 0.05 all trials v.s. random weights, see Methods). (**E**) CoCA projection of neural data in the EPM (top) and rat exposure (bottom) assays for the same mouse. (**F**) Similar to (E), but as a heatmap using testing data from all mice for EPM (top) and rat exposure (bottom). (**G**) Projection of testing neural data is correlated with safety score (EPM, left) and x position (Rat, right) for an example mouse (*r* = Pearson’s correlation coefficient). Larger EPM threat scores correspond to locations more near the extreme end of the open arms. Larger x position values correspond to locations more near the rat. (**H**) Average correlation of projection of neural data with EPM threat score and x position (Rat) differs significantly from 0 (Data are represented as mean ± SEM; *n* = 7 mice; EPM *t* = 10.80; Rat *t* = 6.49, \*\*\**p* < 0.001, one-sample *t*-test).

**Figure 4–figure supplement 1.**
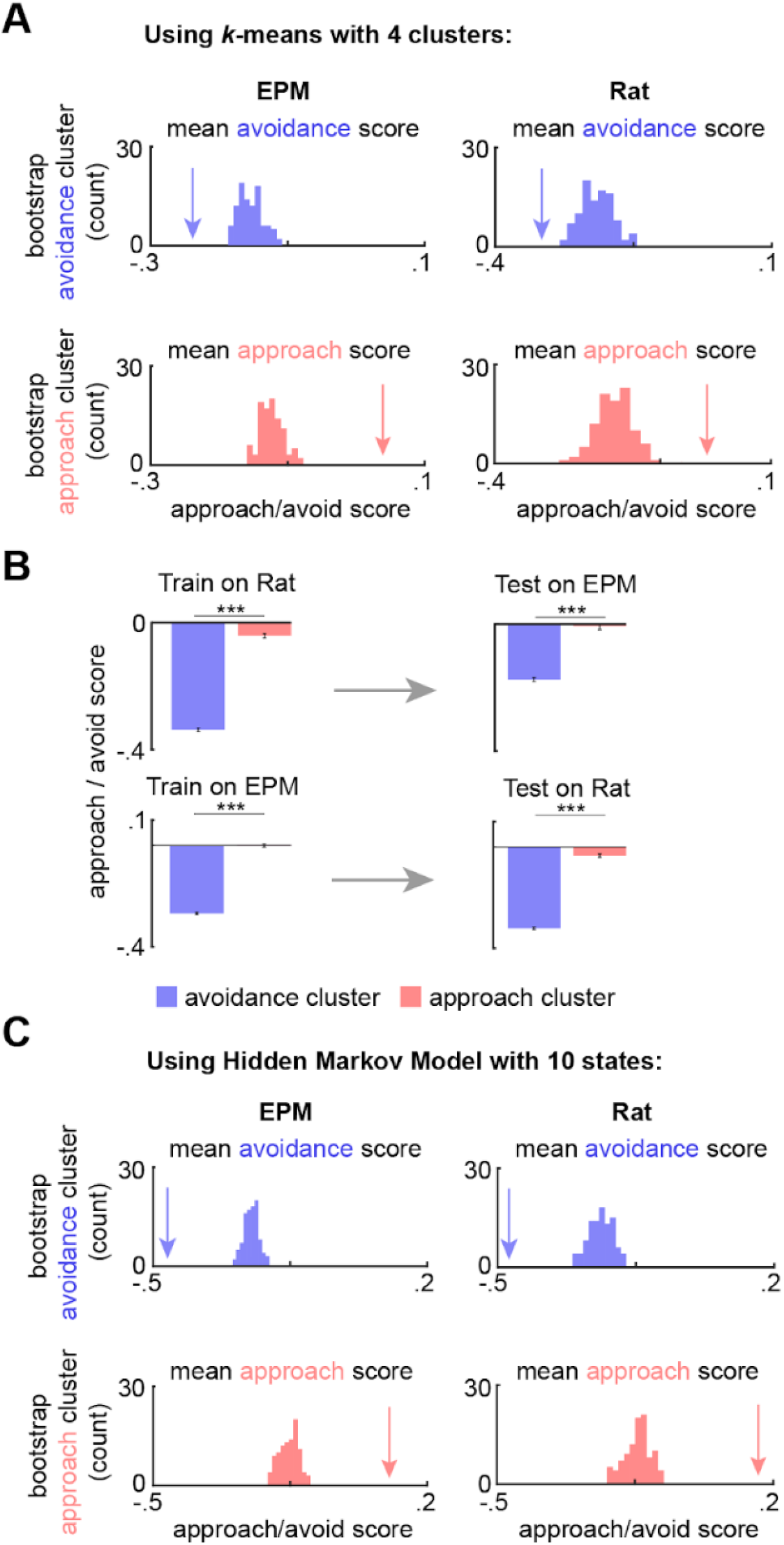
Approach and avoidance identified by k-means with fewer clusters and Hidden Markov Model. (**A**) Arrow depicts the mean approach/avoidance score for avoidance and approach clusters (4 clusters), identified by *k*-means, across concatenated sessions (*n* = 7 mice). This mean was compared to a bootstrapped distribution of approach/avoidance means, calculated by shuffling the neural data 100 times (n cells same as Fig. 4C; *p* < 0.01). (**B**) (left) Bars depict the mean Rat and EPM approach/avoidance scores (+/− 1 s.e.m.) for approach and avoidance clusters across mice. (right) As described in Methods and Fig. 4B, these cluster centroid locations, trained on one assay, were then used to define approach and avoidance timepoints in the other assay. Bars depict the corresponding mean approach/avoidance score (+/− 1 s.e.m.) for this testing data (Train on EPM: avoidance cluster *n* = 14683, approach cluster n=8949; Test on Rat: avoidance cluster *n* = 15727, approach cluster *n* = 7724; Train on Rat: avoidance cluster *n* = 10335, approach cluster *n* = 7742; Test on EPM: avoidance cluster *n* = 9867, approach cluster *n* = 3028; n cells same as Fig. 4D; Wilcoxon ranked-sum test, *** *p* < 0.001). (**C**) Same as (A) but using Hidden Markov Model with 10 states rather than *k*-means; for all, *p* < 0.01).

**Figure 4–figure supplement 2.**
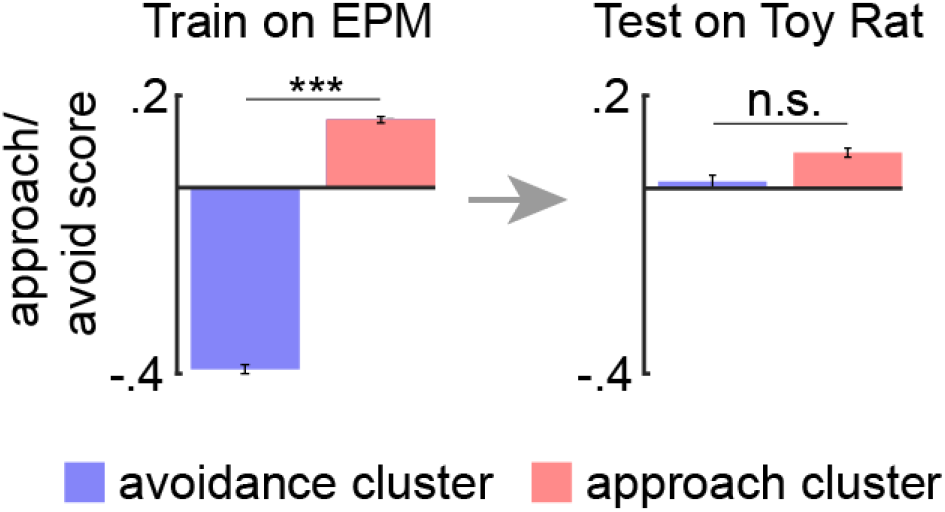
DPAG ensembles to not encode approach and avoidance to a control toy rat. Bars depict the mean EPM approach/avoidance scores (+/− 1 s.e.m.) for approach and avoidance clusters across mice (left). As described in Methods and Figure 4B, these cluster centroid locations, identified using EPM data, were then used to define approach and avoidance timepoints in the toy rat assay (right). Bars depict the corresponding mean approach/avoidance score (+/− 1 s.e.m.) for this testing data (EPM: avoidance cluster *n* = 3066, approach cluster *n* = 6207; Toy Rat: avoidance cluster *n* = 1985, approach cluster *n* = 3497; Wilcoxon ranked-sum test, *** *p* < 0.001, *n* = 7 mice).

**Figure 4–figure supplement 3.**
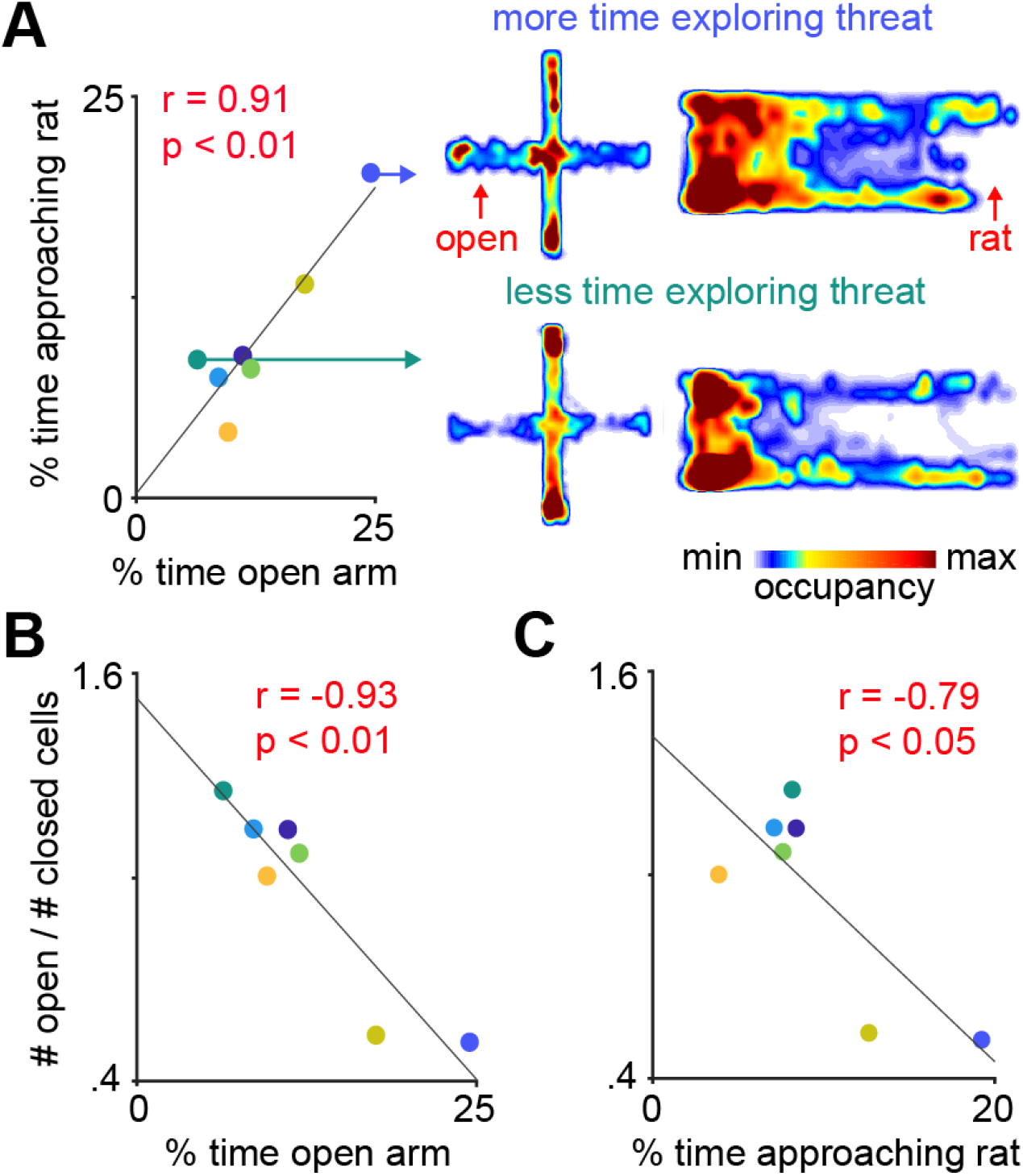
Fraction of open arm cells was negatively correlated with approach to threat across mice. (**A**) Exploration of the open arms in the EPM and approach to rat are correlated. Each point represents one mouse (*n* = 7 mice). Heat maps show exploration in the EPM and the rat assay for example mice showing high (top traces) and low (bottom traces) levels of approach to the open arms and the area near the rat (both high threat areas are shown in red). (**B-C**) Mice with a higher fraction of open arm cells show lower exploration of the open arms in the EPM (**B**) and less time approaching the rat (**C**). *r* = Pearson’s correlation coefficient.

## Materials and Methods

### Mice

Mice (*Mus musculus*) of the C57BL/6J strain (Jackson Laboratory stock No. 000664) were used for all experiments. Male mice between 2 and 5 months of age were used in all experiments. Mice were maintained on a 12-hour reverse light-dark cycle with food and water *ad libitum*. Sample sizes were chosen based on previous behavioral studies with miniaturized microscope recordings on defensive behaviors, which typically use 6-10 mice per group. All mice were handled for a minimum of 5 days prior to any behavioral task. In this work, analyses of the Elevated Plus Maze (EPM) environment used 8 mice, while any analyses involving rat exposure used 7 mice. Sample size was chosen based on prior dlPAG calcium transient recordings (Evans et al., 2018). All procedures have been approved by the University of California, Los Angeles Institutional Animal Care and Use Committee, protocols 2017-011 and 2017-075.

### Rats

Male Long-Evans rats (250-400 grams) were obtained from Charles River and were individually housed on a standard 12-hour light-dark cycle and given food and water *ad libitum*. Rats were only used as a predatory stimulus. Rats were handled for several weeks prior to being used and were screened for low aggression to avoid attacks on mice. No attacks on mice were observed in this experiment.

### Surgeries

Eight-week-old mice were anaesthetized with 1.5-3.0% isoflurane and placed in a stereotaxic apparatus (Kopf Instruments). AAV9.Syn.GCaMP6s.WPRE.SV40 were packaged and supplied by UPenn Vector Core at titers 7.5 × 10^3^ viral particles per ml and viral aliquots were diluted prior to use with artificial cortex buffer to a final titer of 5 × 10^12^ viral particles per ml. After performing a craniotomy, 100nl of virus was injected into the dPAG (coordinates in mm, from skull surface): −4.20 anteromedial, −0.85 lateral, −2.3 depth, 15-degree angle. Five days after virus injection, the animals underwent a second surgery in which two skull screws were inserted and a microendoscope was implanted above the injection site. A 0.5 mm diameter, ~4 mm long gradient refractive index (GRIN) lens (Inscopix, Palo Alto, CA) was implanted above the dPAG (−2.0 mm ventral to the skull surface) (Resendez et al., 2016). The lens was fixed to the skull with cyanoacrylate glue and adhesive cement (Metabond; Parkell). The exposed end of the GRIN lens was protected with transparent Kwik-seal glue and animals were returned to a clean cage. Two weeks later, a small aluminum base plate was cemented onto the animal’s head on top of the previously formed dental cement. Animals were provided with analgesic and anti-inflammatory (carprofen).

### Behavioral timeline

Behavioral tests were combined in the following manner across days: EPM test, habituation 1, habituation 2 (toy rat), rat exposure. Each experiment was performed twice, with two cohorts of 3 and 4 mice each. Each mouse was only exposed to each assay once, as fear assays cannot be repeated. Thus, there are no technical replicates. No outliers were found or excluded. All mice were used. Neural recordings were obtained from all mice in identical conditions, and thus they were all allocated to the same experimental group. There were no experimentally controlled differences across mice and there were no “treatment groups”.

### Elevated Plus Maze test

Mice were placed in the center of the EPM facing one of the closed arms and were allowed to freely explore the environment for 20 minutes. The length of each arm was 30 cm, the width was 7 cm and the height of the closed arm walls was 20 cm. The maze was 65 cm elevated from the floor by a camera stand. A total of 8 mice were analyzed.

### Rat Exposure Assay

Mice were habituated to a white rectangular box (70 cm length, 26 cm width, 44 cm height) for two consecutive days during 20-minute sessions. Mice were then exposed to an adult rat in this environment on the following day. The rat was secured by a harness tied to one of the walls and could freely ambulate only within a short perimeter. The mouse was placed near the wall opposite to the rat and freely explored the context for 20 minutes. No separating barrier was placed between the mouse and the rat, allowing for close naturalistic encounters that can induce a variety of robust defensive behaviors. A total of 7 mice were analyzed.

### Behavior and miniscope video capture

All videos were recorded at 30 frames/sec using a Logitech HD C310 webcam and custom-built head-mounted UCLA miniscope (Aharoni and Hoogland, 2019). Open-source UCLA Miniscope software and hardware (http://miniscope.org/) were used to capture and synchronize neural and behavioral video (Cai et al., 2016; Schuette et al., 2020).

### Perfusion and histological verification

Mice were anesthetized with Fatal-Plus and transcardially perfused with phosphate buffered saline followed by a solution of 4% paraformaldehyde. Extracted brains were stored for 12 hours at 4°C in 4% paraformaldehyde. Brains were then placed in sucrose solution for a minimum of 24 hours. Brains were sectioned in the coronal plane in a cryostat, washed in phosphate buffered saline and mounted on glass slides using PVA-DABCO. Images were acquired using a Keyence BZ-X fluorescence microscope with a 10 or 20X air objective.

Data Analysis was performed using custom-written code in MATLAB and Python.

### Miniscope postprocessing and co-registration

Miniscope videos were motion-corrected using the open-source UCLA miniscope analysis package (https://github.com/daharoni/Miniscope_Analysis) (Aharoni and Hoogland, 2019). They were spatially downsampled by a factor of two and temporally downsampled by a factor of four, and the cell footprints and activity were extracted using the open-source package Constrained Nonnegative Matrix Factorization for microEndoscopic data (CNMF-E; https://github.com/zhoupc/CNMF_E) (Zhou et al., 2018). Neurons were co-registered across sessions using the open-source probabilistic modeling package CellReg (https://github.com/zivlab/CellReg) (Sheintuch et al., 2017).

### Artifact suppression

For suppression of long timescale artifacts, e.g. long-time scale fluctuations in calcium fluorescence shared across many neurons due to bleaching or other factors, we used PCA to identify large variance PCs (≥ 5% total variance) reflecting these artifacts. Cell activity was then reconstructed using these PCs excluded from reconstruction (O’Shea and Shenoy, 2018). This method was applied only to data for mouse 1 in the rat exposure assay.

### Variance thresholding

A minority of recorded cells had very small variance over the course of an experimental session. To exclude these cells from analysis, we identified a representative cell for each trial. Cells with less than 10% of the representative cell’s variance were discarded. The remaining cells were used for further analysis.

### Behavior detection

To extract the pose of freely-behaving mice in the described assays, we implemented DeepLabCut (Mathis et al., 2018), an open-source convolutional neural network-based toolbox, to identify mouse nose, ear and tail base xy-coordinates in each recorded video frame. These coordinates were then used to calculate velocity and position at each time point, as well as classify defensive behaviors in an automated manner using custom Matlab scripts. Freezing was defined as epochs of cessation of all movement except for breathing. Approach and escape were defined as epochs when the mouse moved, respectively towards or away from the rat at a velocity exceeding a minimum threshold.

### Categorization of open, neither and closed arm-preferring cells

A cell was categorized as an open arm-preferring cell if activity in each individual open arm was significantly greater than the pooled activity in the closed arms (Wilcoxon rank sum test, p < 0.05). Likewise, a closed arm-preferring cell was identified as a cell whose activity in each individual closed arm was significantly greater than the pooled activity in the open arms. The remaining cells were labeled as neither arm-preferring cells.

### Behavior-aligned trace and ΔdF/F activity

We calculated each cell’s z-scored behavior-aligned activity by computing the mean activity of the cell over all behavior occurrences, aligned to behavior onset. The mean peri-behavior trace for an ensemble (e.g., closed cells, open cells, or neither cells) was the average of peri-behavior activity across all cells in the ensemble. Change in mean activity after and before behavior was calculated by first subtracting the mean activity of each cell during the time frame [−2.5, 0] seconds relative to behavior onset from the mean activity of each cell in the time frame [0, 2.5] seconds. The overall difference in an ensemble, denoted ΔdF/F, was the average of the change in mean activity across all cells in the ensemble. For head dips, ΔdF/F was calculated using windows of [−5, −2.5] (before) and [0, 2.5] (after).

### Interleaved training and testing data

For analyses involving regression (EPM safety score, CoCA), testing data were interleaved with training data, with 60 seconds for each segment and 10 seconds of separation between data types, i.e., [60s training, 10s excluded, 60s testing, 10s excluded, 60s training, etc.]. These gaps minimize overlapping activity in the training and testing sets, which may arise due to dynamics in calcium activity.

### EPM arm score

The arm score quantifies the separability of cell activity between arm types and is invariant to scaling and shifting. Excluding times when the mouse was in the center of the EPM, data points were labeled according to whether the mouse was in an open arm (positive label) or a closed arm (negative label). The arm score was then defined as:

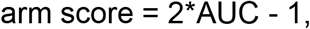

where AUC is the area under the receiver operating characteristic curve resulting from predicting which arm the mouse was in from single-cell activity. Cells with strong preference for firing in the closed arms or open arms respectively have arm score values near −1 and +1. An arm score of exactly +1 indicates that cell activity in the open arm is strictly greater than activity in the closed arm.

### EPM score

The EPM score quantifies how differently a cell fires between closed vs open arms. It is close to 1 when the cell has large differences in activity between arms of different types but is negative if the cell’s activity is more similar between different arm types than between same arm types. To calculate EPM score, we first compute the mean difference in z-scored activity between arms of different types (A) and arms of the same type (B). These are defined as

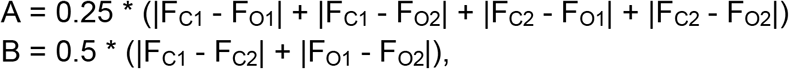

where F_O1_ and F_O2_ are the mean z-scored activity of the cell in open arms 1 and 2, respectively, and F_C1_ and F_C2_ are the mean z-scored activity in closed arms 1 and 2 (Adhikari et al., 2011). The EPM score is defined as:

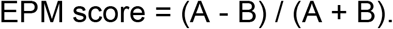

Cells with high EPM score if they have large differences in activity in different arm types (large A) and similar activity in same type arms (small B). The maximum score of 1.0 indicates no difference in firing rates across arms of the same type (B = 0). Cells with negative EPM scores have more similar activity across arms of different types than across arms of the same type.

### EPM threat score

The threat score quantifies the threat exposure to the mouse. It is close to 1 when the mouse is at the end of an open arm, and close to −1 when the mouse is at the end of the closed arms. To calculate threat score, we first normalized the x and y position of the mice in the EPM to be in the range [−1, 1], where the x position is ±1 at the ends of the open arms and the y position is ±1 at the ends of the closed arms. We defined the threat measure as:

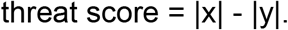

The threat score is therefore a value between-1 and 1. Prediction of threat score (Figure 1H-I) was performed using linear regression with interleaved training and testing data. Outputs were clipped to the range [−1, 1] before final prediction was made.

### Prediction of mouse position in the EPM from neural data using a linear support vector machine (SVM)

After z-scoring data, times when the mouse was in the center of the EPM were removed from training. The remaining data were separated into alternating blocks of 50s training data and 50s testing data with 10s of separation between blocks. A linear SVM was fit on training using scikit-learn’s SVC function with balanced class weights (Pedregosa et al., 2011). Significance testing was performed with bootstrapping using shuffled class labels for 100 random trials per mouse. The Matthews correlation coefficient was used to quantify the relation between predicted and observed arm-type occupancy because this metric was developed to assess correlations between binary values (such as arm-type, which can only be closed or open arms).

### Zones in the rat assay

The safe zone was defined as the left 20% of the rat environment, based on x position. The threatening zone was defined as being within 20% of the maximum distance from the rat during the exposure.

### Generalized linear model

A generalized linear model (GLM) was fit for each cell. Each GLM mapped behavior variables to the cell’s z-scored calcium activity. Discrete behaviors were binary, labeled as 1 at all times in which they occurred and 0 otherwise. To enable behaviors to alter neural activity prior to and following the behavior, each binarized behavior was convolved with a non-causal log-time scaled raised cosine basis, from 5 seconds before behavior onset to 5 seconds after behavior offset. Further, to enable historical kinematics to affect present neural data, the kinematics were convolved with a causal kernel, which used the same set of bases as the behavior, but only had responses after the onset within 5 seconds. These convolved behavior variables, denoted here as y_1_, y_2_, etc., were then modeled to produce the cell’s calcium fluorescence as:

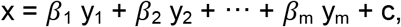

where *β*_i_ is the coefficient for the ith behavior variable. In total, there were 8 behavior variables for rat: distance to rat, mouse velocity, rat velocity, angle from mouse’s head to the rat, and occurrence of approaching, escaping and freezing. The GLM was optimized by minimizing the mean-square error of the reconstruction between the GLM activity estimate, x, and the recorded calcium activity.

### Cross-assay Constrained Correlation Analysis (CoCA)

To investigate if the dPAG uses shared patterns of neural activity to represent threat imminence across assays we developed Constrained Correlation Analysis (CoCA). This model identifies the features used in a shared neural representation between different threatening situations. The CoCA technique defined a shared neural projection as a linear combination of the activity of individual neurons. Additionally, we constructed a behavioral projection for each assay, through a linear combination of each assay’s behavioral variables. Optimization via CoCA produced neural projection weights that were compared across open and closed cell ensembles.

We denote calcium fluorescence neural data as *X* ∈ R^kxT^ and externally-observed behavioral data as *Y* ∈ R ^pxT^, where k is the number of recorded cells for the corresponding mouse, shared across assays, p is the chosen number of behavioral variables for the corresponding assay, shared across mice, and T is the length of a recording session, unique for each session, but shared between neural and behavioral data for the same session. Behavioral variables contained both continuous kinematic variables (such as speed and distance from rat) as well as binary defensive behavior variables (such as the occurrence of freezing and escape). All variables were normalized to zero mean and unit variance, except normalized |x| and |y| in EPM, which were already in the range [0, 1] (variance < 0.25).

In order to find a common linear projection of threat across mice and assays, we performed the following optimization with mouse IDs *i = 1, 2,…;7* and assay IDs *j = EPM, RAT*. Calcium fluorescence traces of dPAG cells for mouse *i* were linearly combined after multiplying each cell with weights n_1_^i^ to n_k_^i^, where k is the number of cells that were co-registered in both assays. Taking the dot product of the calcium activity for mouse *i* in assay *j*, given by X_i,j_, and the weights **n**^i^ = [n^i^_1…k_] defined a neural projection for mouse *i* and assay *j*, given by N_i,j_ = (**n**^i^)^T^X_i,j_ (Figure 6A, neural data projection in red). For each mouse, the weights, **n**^i^, were the same across assays, so that each cell had the same weight in both assays. The behavioral variables for the EPM (such as x and y position, speed, etc.) were linearly combined with a set of weights b_1_ to b_6_ (as 6 behavioral variables were used for the EPM). These weights, **b**^EPM^ = [b_1_^EPM^, b_2_^EPM^,…;, b_6_^EPM^] were conserved across all mice. Linearly combining the EPM behavioral variables resulted in a behavioral projection for mouse *i* and assay *EPM*, given by B_i,EPM_ = (**b**^EPM^)^T^Y_i,EPM_. Similarly, 9 behavioral variables from the rat assay were linearly combined to produce a behavioral projection B_i,RAT_ = (**b**^RAT^)^T^Y_i,RAT_ using weights **b**^RAT^ = [b_1_ ^RAT^, b ^RAT^,…;, b ^RAT^]. We chose the neural weights, **n**^i^, and the behavioral weights, **b**^EPM^ and **b**^RAT^, to optimize the correlations across all mice and assays:

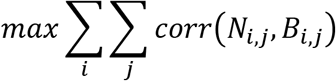

where corr() is the Pearson correlation coefficient, and N_i,j_ and B_i,j_ are the linear projections of neural and behavioral data, respectively, given by: N_i,j_ = (**n**^i^)^T^X_i,j_ and B_i,j_ = (**b**^j^)^T^Y_i,j_. The optimization variables **n**^i^, *i = 1, 2,…;7*, and **b**^j^, *j = EPM, RAT*, were simultaneously optimized using gradient descent via the Adam optimizer (Kingma and Ba, 2015) until convergence. Results presented use interleaved training and testing data. This method was implemented using PyTorch.

### CoCA bootstrapping

In order to test if correlations of testing data were better than expected by chance, correlations were computed between projected behavioral data (using projections fit by training data) and random projections of neural data (1000 trials). We emphasize these correlations were applied to the testing data, and therefore it was possible for a random projection to have higher correlation than the CoCA projection. Here a one-tailed test was used.

### Approach/Avoidance score

To calculate the continuous approach/avoidance score for each assay, the distance from safety was calculated (Rat: distance from the safe wall; EPM: distance from the end of the closed arm) and normalized such that it ranged from 0 to 1 in the Rat assay and 0 to 0.9 in the EPM. A binarized direction value was also assigned to each timepoint, indicating if the mouse was moving towards (+1) or away from (−1) the threat. To incorporate categorized behaviors, the approach/avoidance score for freeze samples equaled the minimum score of −1. For the EPM only, the approach/avoidance score was multiplied by 1.11 for head dip samples, such that a head dip at the end of the open arm would yield the maximum score of 1.

To calculate the score at each timepoint:

While approaching threat, approach/avoidance score = distance to safety x direction

While avoiding threat: approach/avoidance score = [1-distance to safety] x direction

### K-means clustering of neural data

To determine if the approach/avoidance score is represented in the neural data, the k-means algorithm (k=10) was used to cluster the neural data in an unsupervised manner. For each implementation of k-means, ten sets of clusters were identified using ten different randomized initializations; the set with the minimum sum of euclidean distances was used. The approach and avoidance clusters then identified, for each session, as those with, respectively, the highest and lowest mean approach/avoidance scores. The overall mean approach/avoidance scores for approach and avoidance clusters were then calculated across mice. To determine if these approach and avoidance cluster scores were statistically significant, the actual mean was compared to a bootstrapped distribution of means, calculated in an identical manner with shuffled neural data over 100 iterations. If the approach and avoidance score means were respectively greater than or less than 95% of this bootstrapped distribution, they were considered significant. For the training/testing analysis, *k*-means was implemented on one assay as described above (the training assay), using only cells that coregistered between both assays. The cluster centroids identified in the training assay were then used to categorize approach and avoidance samples in the withheld testing assay. The mean approach/avoidance score was calculated for all approach and avoidance timepoints, across all training or testing sessions.

In a similar way, approach/avoidance states were identified by Hidden Markov Models (10 states), using the top principal components of the neural data as input (accounting for >=60% of the total variance). These states were analyzed in an identical manner to the k-means clusters described above. For the code, see ‘Expectation-Maximization for Hidden Markov Models using real-values Gaussian observations’ at Zoubin Ghahramani’s code base: http://mlg.eng.cam.ac.uk/zoubin/software.html).

### Statistical analysis

Significance values are included in the figure legends. Unless otherwise noted, all statistical comparisons were performed by either nonparametric Wilcoxon rank-sum or signed-rank tests. With the exception of CoCA bootstrapping, all significance tests were two-tailed. Standard error of the mean was plotted in each figure as an estimate of variation of the mean. Correlations were calculated using Pearson’s method. Multiple comparisons were corrected with the false discovery rate method. All statistical analyses were performed using SciPy (Virtanen et al., 2020) and custom Matlab scripts.

### Data and code availability

All data was uploaded to dryad and all code was uploaded to github.

https://datadryad.org/stash/share/4GezSjw4dvDJClAWa_zRoNWioH9qzGtDCJjLQ89HVoA

https://doi.org/10.5068/D1TM2G

https://github.com/schuettepeter/eLife_dPAG-ensembles-represent-approach-and-avoidance-states

